# Autogenous cross-regulation of *Quaking* mRNA processing and translation balances *Quaking* functions in splicing and translation

**DOI:** 10.1101/127936

**Authors:** W. Samuel Fagg, Naiyou Liu, Jeffrey Haskell Fair, Lily Shiue, Sol Katzmann, John Paul Donohue, Manuel Ares

## Abstract

Quaking protein isoforms arise from a single *Quaking* gene, and bind the same RNA motif to regulate splicing, translation, decay, and localization of a large set of RNAs. However, the mechanisms by which *Quaking* expression is controlled to ensure that appropriate amounts of each isoform are available for such disparate gene expression processes are unknown. Here we explore how levels of two isoforms, nuclear Quaking-5 (Qk5) and cytoplasmic Quaking-6 (Qk6), are regulated in mouse myoblasts. We find that Qk5 and Qk6 proteins have distinct functions in splicing and translation respectively, enforced through differential subcellular localization. We show that Qk5 and Qk6 regulate distinct target mRNAs in the cell, and act in distinct ways on their own and each other’s transcripts to create a network of auto- and cross-regulatory feedback controls. Morpholino-mediated inhibition of Qk translation confirms that Qk5 controls *Qk* RNA levels by promoting accumulation and alternative splicing of *Qk* RNA, whereas Qk6 promotes its own translation while repressing Qk5. This Qk isoform cross-regulatory network responds to additional cell type and developmental controls to generate a spectrum of Qk5/Qk6 ratios, where they likely contribute to the wide range of functions of *Quaking* in development and cancer.

## Introduction

Sequence-specific RNA-binding proteins (RBPs) regulate nearly every posttranscriptional step of gene expression. Recognition of cis-acting regulatory sequences by the RNA-binding domain (RBD), is followed by one or more consequences, including RNA refolding, folding or refolding of the protein itself, leading to altered interactions and functional impact on RNA polymerase, the spliceosome, the export and decay machineries, and the ribosome (Keene 2007; Fu and Ares 2014). Furthermore, many RBPs control their own levels (autogenous regulation), through the same mechanisms by which they regulate their targets (Wollerton et al. 2004; Lareau et al. 2007; Ni et al. 2007; Damianov and Black 2010; Sun et al. 2010). In many metazoans, multiple RBPs with the same or highly similar sequence binding properties are present at the same time in the same cells but perform distinct functions (Boutz PL 2007a; Makeyev et al. 2007; Spellman et al. 2007; Yeo et al. 2009; Gehman et al. 2011; Charizanis et al. 2012; Gehman et al. 2012; Singh et al. 2014), suggesting that functional diversification is enforced by regulatory mechanisms that prevent crosstalk. These two circumstances, autogenous regulation through binding their own pre-mRNAs and mRNAs, combined with functional diversification using the same recognition sequences, create a regulatory conundrum: How is the right amount of each distinctly functional RBP isoform established and maintained in cells?

Since all isoforms within a family recognize essentially the same RNA sequence, control of individual isoforms is unlikely to be mediated solely through their common RNA target sequences. Additional mechanisms for distinguishing isoforms include 1) isoform-specific localization that restricts function to processes within specific compartments (Caceres et al. 1998; Koizumi et al. 1999; Cazalla et al. 2002; Sanford et al. 2004), and 2) isoform-specific protein-protein interactions with different core machineries (e. g. spliceosome, ribosome) (Roscigno and Garcia-Blanco 1995; Jin et al. 2004; Cheever and Ceman 2009; Sharma et al. 2011). Because many RBPs are encoded by multi-gene families with complex alternative splicing (Underwood et al. 2005), whole menageries of functionally distinct RBP isoforms that recognize the same RNA target can be expressed within single cell types (Lee et al. 2009; Hamada et al. 2013; Lee et al. 2016). How does the cell recognize when each distinctly functional RBP family member is present in an appropriate amount for each process?

To examine this question, we have focused on the single *Quaking* gene (*QKI* in human, *Qk* in mouse), which is required for a broad set of functions in diverse tissues (Ebersole et al. 1996; Zhao et al. 2010; Darbelli et al. 2016; de Bruin et al. 2016), through its contribution to RNA processing steps including splicing (Hall et al. 2013; van der Veer et al. 2013; Darbelli et al. 2016), localization (Li et al. 2000; Larocque et al. 2002), stability/decay (Li et al. 2000; Larocque et al. 2005; Zearfoss et al. 2011; de Bruin et al. 2016), translation (Saccomanno L 1999; Zhao et al. 2010), and miRNA processing (Wang et al. 2013; Zong et al. 2014). These processes are regulated by dimeric Qk binding an RNA element that includes ACUAAY and a “half-site” (UAAY) separated by at least one nucleotide (Ryder and Williamson 2004; Galarneau and Richard 2005; Beuck et al. 2012; Teplova et al. 2013). *Quaking* gene transcription initiates primarily at a single major site, and in most cell types, three alternatively spliced mRNAs encode three protein isoforms [Quaking-5 (Qk5), Quaking-6 (Qk6), and Quaking-7 (Qk7)] that differ only in the C-terminal tail (Ebersole et al. 1996; Kondo et al. 1999). Although various cell types express different ratios of Qk protein isoforms (Ebersole et al. 1996; Hardy et al. 1996; Hardy 1998; van der Veer et al. 2013; de Bruin et al. 2016), it is unclear how the relative isoform ratios are maintained in order to support tissue-specific regulated RNA processing. Disruption of these ratios is associated with developmental defects (Ebersole et al. 1996; Cox et al. 1999), cancer (de Miguel et al. 2016; Sebestyen et al. 2016), and schizophrenia (Aberg et al. 2006).

Many studies of *Quaking* function have used overexpression of Qk isoforms (Wu et al. 2002; Hafner et al. 2010; Wang et al. 2013), or depletion strategies and mutant models that do not distinguish which Qk isoform is functional (Hardy et al. 1996; Lu et al. 2003; van der Veer et al. 2013; Darbelli et al. 2016). Here we test specific Qk isoforms for separate functions, and identify in part how the appropriate balance of Qk isoforms is maintained. In mouse myoblasts, Qk5 and Qk6 are the predominantly expressed isoforms and we find that Qk5, but not Qk6, regulates splicing, while Qk6 controls mRNA translation and decay. This functional specificity is mediated by subcellular localization encoded into the unique C-terminal amino acids of these isoforms. Furthermore, the relative expression of Qk protein isoforms is regulated in part, by Qk protein isoforms themselves through both auto- and cross-regulatory influences characteristic of the function of each isoform on its other RNA targets. These findings uncover unexpectedly complex isoform control within a single family of RBPs, and suggest that the relative amounts of each isoform are set in a cell type specific fashion and homeostatically controlled by Qk protein isoform levels themselves.

## Results

### Qk5 and Qk6 are the predominant isoforms in myoblasts

We analyzed the abundance and localization of Qk isoforms (Fig 1A) in myoblasts and differentiated myotubes (Yaffe and Saxel 1977) using isoform-specific antibodies. Total Qk protein level increases during C2C12 myoblast differentiation ((Hall et al. 2013), Fig 1B), with Qk5 the most abundant, followed by Qk6, and then Qk7 (Fig 1B). During differentiation each isoform increases proportionately (Fig 1B) and total Qk protein remains predominantly localized in nuclei (Fig S1A). Immunolocalization using isoform-specific antibodies shows that Qk5 is primarily nuclear, although some cytoplasmic localization is observed, whereas Qk6 and Qk7 are present in both the nuclear and cytoplasmic compartments (Fig 1C). Cell to cell heterogeneity observed for nuclear Qk6 and Qk7 was sometimes evident (Fig 1C), but was judged to be minor after quantification of nuclear/cytoplasmic ratios for many cells by high throughput image analysis (Fig 1D). Although the precise ratios using this two-dimensional method are subject to cytoplasmic signal that overlays the nucleus, we conclude that the major isoforms in myoblasts are distributed in distinct nuclear/cytoplasmic ratios, with Qk5 being predominantly nuclear, and with Qk6 and Qk7 distributed throughout cells, but with Qk6 being more cytoplasmic than Qk7.

**Figure 1:**
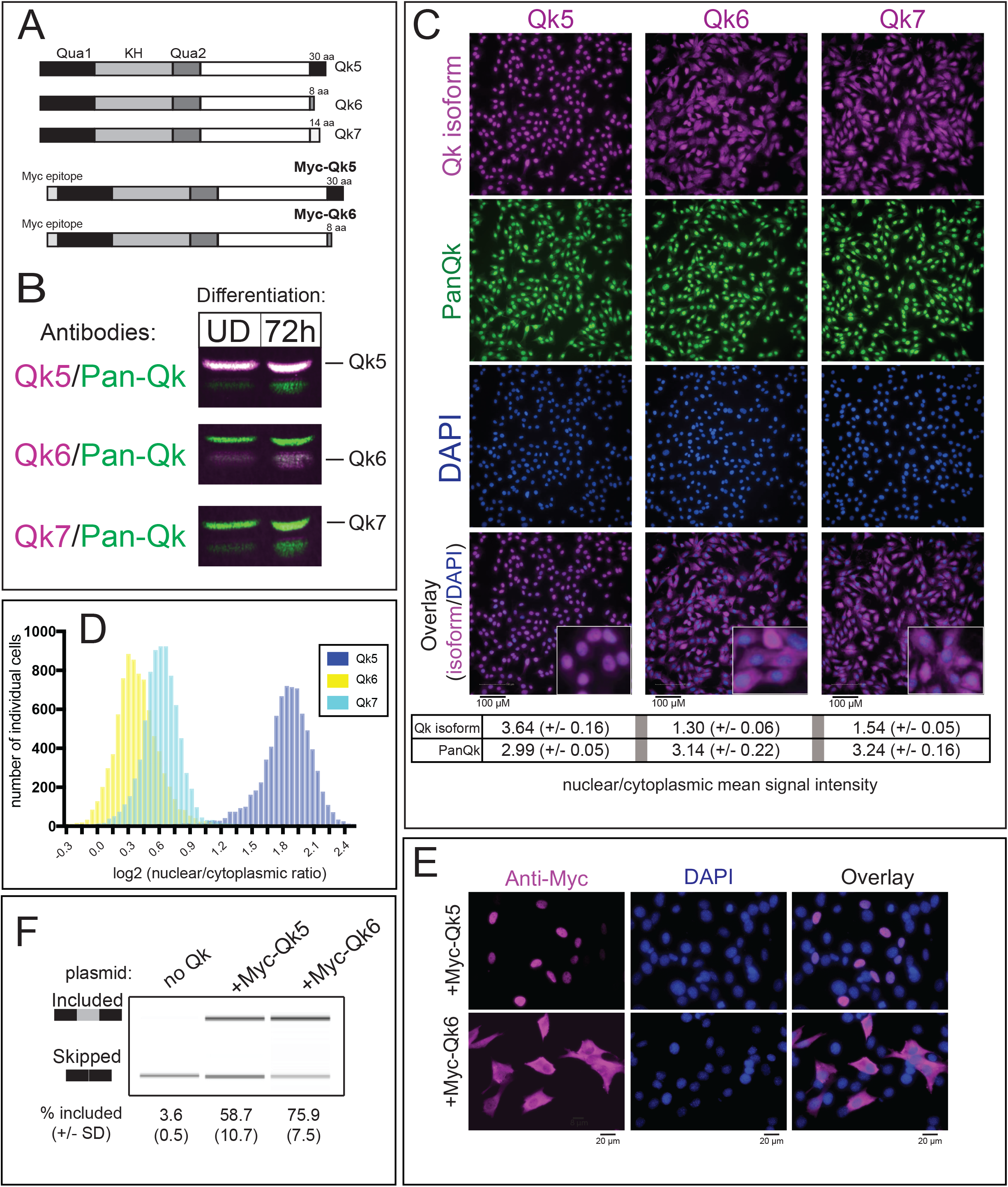
Qk5 and Qk6 are the predominant isoforms and can activate splicing in C2C12 muscle cells A. Model depicting endogenous Qk protein isoforms (top) with number of unique amino acids shown at C-terminus (right), and Myc-epitope tagged constructs (bottom) used in subsequent experiments; all Qk proteins share the same 311 amino acids including Quaking homology 1 (Qua1) and 2 (Qua2), and hnRNP-K homology (KH) domains. B. Western blot of whole cell extracts prepared from undifferentiated C2C12 myoblasts (UD) or C2C12 cells differentiated for 72h (72h) simultaneously probed with infrared-conjugated secondary antibodies to Qk5 (top), Qk6 (middle), or Qk7 (bottom) in magenta (notation to the right indicates migration of specific isoform) or PanQk in green, which recognizes all Qk isoforms. C. Indirect immunofluorescence showing Qk isoform-specific (magenta), PanQk (green), and DAPI (blue) staining in C2C12 myoblasts with overlay of Qk isoform and DAPI at bottom with higher magnification inset; mean signal intensity for each Qk isoform and total Qk (PanQk) is shown at bottom (+/- standard deviation) calculated for total well (n = 9; see methods section for details); scale bar represents 100 µm. D. Histrogram of log_2_ nuclear/cytoplasmic signal intensity of Qk isoforms (x-axis) calculated for each cell imaged in C., with number of cells shown on y-axis. E. Indirect immunofluorescence of C2C12 myoblasts transfected with either Myc-Qk5 (top) or Myc-Qk6 (bottom) stained using Anti-Myc epitope antibody in magenta (left), DAPI in blue (middle), and both channels overlaid (right); scale bar represents 20 µm. F. RT-PCR products from Dup-Capzb exon 9 splicing reporter analyzed on BioAnalyzer from RNA extracted from C2C12 myoblasts either mock transfected, or transfected with Myc-Qk5 or Myc-Qk6 plasmid; mean values of percent included are shown below (+/- standard deviation) from three independent biological replicates.

### Qk-dependent alternative splicing regulation requires Qk5, but not Qk6

We previously showed that depletion of all Qk isoforms alters splicing of several hundred exons in myoblasts through Qk-responsive ACUAA sequence elements (Hall et al. 2013). To test which isoform is responsible for splicing regulation, we overexpressed a Myc-tagged cDNA construct for each isoform (Fig S1B), and measured splicing of the Qk-regulated Capzb exon 9 in a β–globin reporter (Dominski and Kole 1991; Hall et al. 2013). To our surprise, both isoforms efficiently activate splicing (Fig 1F). Previous work shows that Qk proteins are only stable as a dimer *in vivo* (Beuck et al. 2012), and that overexpression of Qk isoforms can create ectopic Qk heterodimers (Pilotte et al. 2001). Consistent with this, we observe nuclear staining of Myc-Qk6 (Fig 1E). To rigorously test isoform function in splicing, we used a depletion-replacement strategy in which endogenous Qk mRNAs and proteins are depleted using an siRNA to a common Qk region, followed by expression of a single siRNA resistant isoform (kind gift from Sean Ryder), to create a cellular pool of Qk dominated by a single isoform (Fig 2A, Fig S2A). Under the depletion-replacement treatment, inclusion of Capzb reporter exon is specifically increased by replacement with Myc-Qk5, but not with Myc-Qk6 (Fig 2B). In addition, whereas reconstituted Myc-Qk5 accumulates efficiently in the nucleus, Myc-Qk6 remains predominantly cytoplasmic (Fig 2C). Taken together, we conclude that Qk5 is predominantly nuclear and serves as a splicing factor, but Qk6 is not, consistent with this isoform regulating translation (Saccomanno L 1999). We suggest that only when overexpressed Qk6 can create Qk5-Qk6 heterodimers that go to the nucleus via the Qk5 subunit, can Qk6 contribute to splicing regulation. This predicts that isoform function might be enforced by localization.

**Figure 2:**
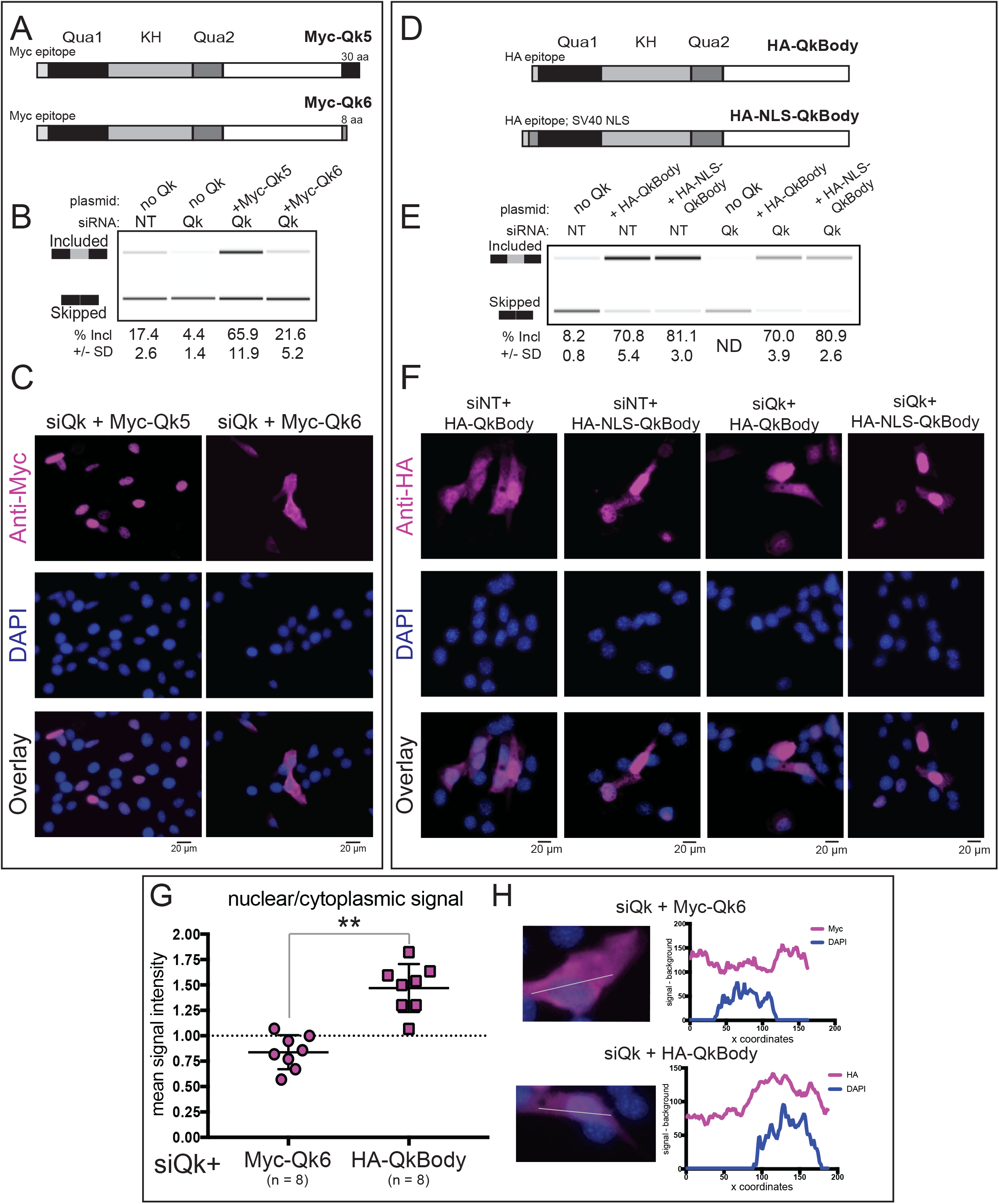
Nuclear localization is sufficient for Qk splicing function. A. Model of Myc epitope-tagged Qk5 and Qk6 with number of unique amino acids shown at C-terminus. B. RT-PCR products from Dup-Capzb exon 9 splicing reporter analyzed on BioAnalyzer from RNA extracted from C2C12 myoblasts transfected with either control siRNA (siNT) or siRNA targeting all Qk isoforms (siQk), or co-transfected with both siQk and Myc-Qk5 or siQk and Myc-Qk6 (top); mean percent included is shown below with standard deviation from three independent biological replicates. C. Indirect immunofluorescence using Anti-Myc epitope antibody in magenta (top), DAPI in blue (middle), and channels overlaid (bottom) of C2C12 myoblasts transfected with siQk + Myc-Qk5 (left) or siQk + Myc-Qk6 (right); scale bar represents 20 µm. D. Model of Qk mutant proteins tested for splicing function and localization: HA-QkBody consists of an N-terminal HA epitope tag and the 311 amino acids common to all Qk proteins (top) and HA-NLS-QkBody consists of an N-terminal HA tag and NLS from SV40 preceding the common 311 amino acids to all Qk isoforms (bottom). E. RT-PCR products from Capzb exon 9 splicing reporter analyzed on BioAnalyzer from RNA extracted from C2C12 myoblasts transfected with siNT, siNT and HA-QkBody, siNT and HA-NLS-QkBody, siQk, siQk and HA-QkBody, or siQk and HA-NLS-QkBody (left to right); mean percent included from three independent biological replicates with standard deviation shown below. F. Indirect immunofluorescence using HA antibody (top, magenta), DAPI staining (blue, middle), and overlaid images at bottom of representative C2C12 myoblasts from siNT + HA-QkBody, siNT + HA-NLS-QkBody, siQk + HA-QkBody, and siQk + HA-NLS-QkBody transfections (left to right); scale bar represents 20 µm. G. Graph showing nuclear/cytoplasmic signal intensity of Myc-Qk6 (left) or HA-QkBody (right) staining of C2C12 myoblasts transfected with siRNA targeting all Qk isoforms (siQk; n = 8; **p < 0.01). H. Representative images (left) from which epitope tag and DAPI signal intensity were measured for data shown in G.; plots on right show corresponding signal intensities (minus background; y-axis) plotted over image coordinates (x-axis) based on the line drawn through representative images on left.

### Nuclear presence of the common Qk protein sequence is sufficient for splicing function

If localization in the nucleus of the common (ie non-isoform specific) parts of Qk is sufficient to activate splicing, then localization of just the common Qk sequences by other means should also serve. The C-terminal 30 amino acids of Qk5 encode a non-canonical NLS (Wu et al. 1999), however this domain could be required for other function(s) specifically necessary for splicing. To test this, we made two Qk constructs lacking any isoform-specific C-terminal amino acids comprising only the common or “body” 311 amino acids shared by all natural Qk isoforms (Fig 2D). One construct encoded an N-terminal HA epitope tag (HA-QkBody), and the other an N-terminal HA tag followed by the canonical SV40 NLS (HA-NLS-QkBody). Upon expression in myoblasts (Fig S2B), these mutant Qk proteins promote inclusion of Capzb exon 9, whether endogenous Qk protein is depleted or not (Fig 2E). This shows that all necessary residues required for Qk splicing functions reside in the body of the Qk protein and not in the Qk5 isoform-specific tail. The HA-QkBody protein is distributed throughout the cell, whereas the HA-NLS-QkBody protein is concentrated in nuclei (Fig 2F), reinforcing the conclusion that nuclear localization is sufficient for splicing function, mediated by the non-canonical NLS-containing Qk5-specific tail. By comparing the subcellular distribution of the HA-QkBody protein to that of Myc-Qk6 in this reconstituted cell system, the contribution of the Qk6 isoform specific tail to cytoplasmic localization can be measured (Fig 2G and 2H). This result indicates that the isoform-specific Qk6 tail contains a cytoplasmic retention or nuclear export signal that limits its contributions to splicing and directs it to cytoplasmic functions like translation, decay and localization.

### Qk5 is required for expression of Qk6 and Qk7

To refine the understanding of Qk isoform-specific functions obtained from cells expressing single Qk protein isoforms, we selectively targeted individual protein isoforms using isoform-specific siRNAs (siQk5 or siQk6/7; see Supplemental Methods). Treatment with siQk5, produces the expected alternative splicing changes (Hall et al. 2013) in the endogenous Rai14 cassette exon 11 (loss of repression, Fig 3A) and the Capzb exon 9 splicing reporter (loss of activation, Fig 3B), whereas treatment with siQk6/7 causes slight but statistically significant responses in the opposite direction. Analysis of the samples for protein depletion reveals that treatment with siQk5 unexpectedly depletes all Qk protein isoforms (Fig 3C), whereas treatment with siQk6/7 leads only to the loss of Qk6 and Qk7 (Figs 3C and S3A). We specifically targeted Qk5 mRNA with two additional siRNAs leading to the same loss of all Qk proteins (Fig S3B). Depletion of Qk6 and Qk7 proteins by Qk5-specific siRNA is as efficient as that observed with a Qk6/7-specific siRNA (Fig S3A and S3B). We interpret these data to mean that cells cannot express Qk6 or Qk7 RNA or protein unless they also or first express Qk5 protein. We conclude that Qk5 is required to promote efficient expression of all the major proteins produced by the *Qk* gene.

**Figure 3:**
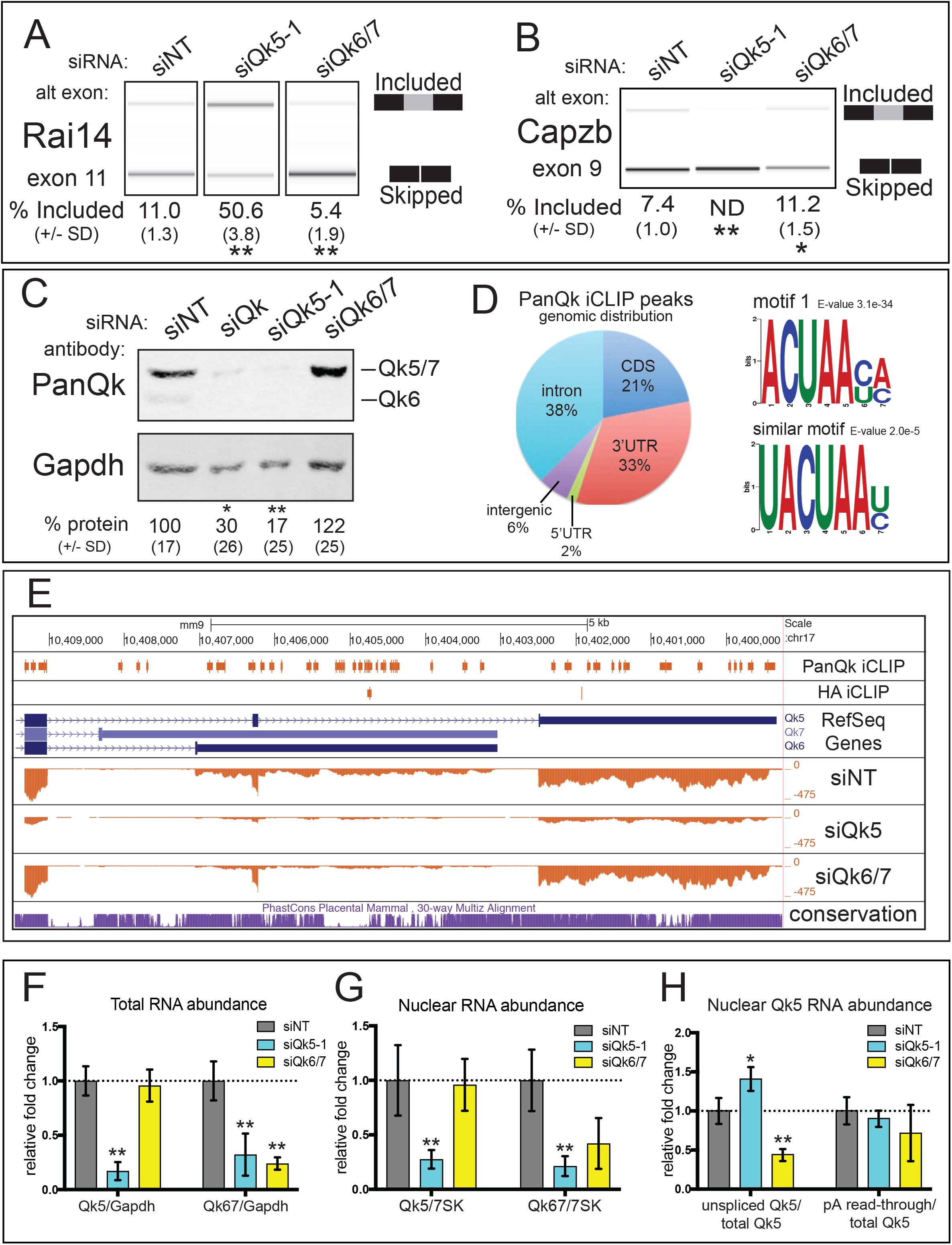
Qk5 is required for splicing and Qk5 and Qk6 accumulation. A. RT-PCR products from endogenous Rai14 exon 11 analyzed on BioAnalyzer from RNA extracted from C2C12 myoblasts transfected with siNT, siQk5-1, and siQk6/7; mean percent included from 3 biological replicates +/- standard deviation is shown below (** p < 0.01). B. RT-PCR products from Dup-Capzb exon 9 splicing reporter analyzed on BioAnalyzer from RNA extracted from C2C12 myoblasts transfected with siNT, siQk5-1, and siQk6/7; mean percent included from 3 biological replicates +/- standard deviation is shown below (** p < 0.01; * p < 0.05). C. Representative western blot of protein extracted from C2C12 myoblasts transfected with siNT, siQk, siQk5-1, and siQk6/7 probed simultaneously with anti-PanQk and anti-Gapdh (notation at right shows migration pattern of Qk protein isoforms); mean relative percentage protein abundance from 3 biological replicates shown below +/- standard deviation (** p < 0.01; * p < 0.05). D. Percentage of iCLIP peaks for PanQk that did not overlap with control HA peaks shown mapped to genome region (left) and motifs found from unique PanQk iCLIP peaks (right). E. UCSC Genome Browser screen shot showing Qk 3’end with PanQk and HA iCLIP peaks, RefSeq genes, coverage tracks for siNT, siQk5-1, and siQk6/7, and conservation (from top to bottom). F. RT-qPCR analysis of RNA extracted from C2C12 myoblast cultures transfected with siNT, siQk5-1or siQk6/7 for Qk5 (left) or Qk6/7 (right) RNA normalized to Gapdh RNA and displayed as fold change relative to siNT; error bars show standard deviation from the mean using three independent biological replicates (** p < 0.01). G. RT-qPCR analysis of nuclear RNA extracted from C2C12 myoblasts cultures transfected with siNT, siQk5-1 or siQk6/7 for Qk5 (left) or Qk6/7 (right) RNA normalized to 7SK RNA and displayed as fold change relative to siNT; error bars show standard deviation from the mean using three independent biological replicates (** p < 0.01). H. RT-qPCR analysis of samples described in G. but measuring unspliced Qk5 or non-poly adenylated Qk5 normalized to total Qk5 and displated as fold change relative to siNT; error bars show standard deviation from the mean using three independent biological replicates (** p < 0.01; p < 0.05).

### Qk5 and Qk6 are required for expression of distinct sets of mRNAs enriched for Qk recognition sequences

The results above suggest at a minimum that Qk5 expression is required for expression of Qk6, probably through its RNA-binding activity. To explore the impact of loss of Qk5 and Qk6 broadly on the myoblast transcriptome, and to map Qk binding to its RNA targets, we performed RNAseq on cells treated with siQk5-1, siQk6/7, or with the non-targeting siNT, and also performed crosslinking immunoprecipitation sequencing (iCLIP (Huppertz et al. 2014)) on unperturbed myoblasts. We mapped and compared CLIP peaks generated using the pan-Qk antibody that recognizes all Qk isoforms, and an HA-antibody control (Supplemental Table 1). Qk-specific peaks indicate that Qk binds mostly introns and 3’ UTRs in myoblasts, and less frequently in coding regions (Fig 3D). The top motif identified in the Pan-Qk CLIP peaks is related to the known Qk responsive element (Fig 3D). Consistent with the idea of cross isoform regulation, and with previous observations in human cell lines, the distinct 3’UTRs of the Qk5 and Qk6 mRNAs both bind Qk protein at numerous sites (Fig 3E).

RNAseq libraries were mapped to the mouse genome (mm9) and analyzed by DESEQ2 (Love et al. 2014) to extract gene specific changes due to each isoform (See Supplemental Methods, and Supplemental Table 2). We identified two sets of genes, one whose mRNAs change significantly up or down upon loss of Qk5, and another whose mRNAs change significantly upon loss of Qk6. Although there is no overlap between these two gene sets (Suppl. Table 2G), we did observe genes whose expression changed significantly in different directions when Qk5 (and Qk6) are depleted by siQk5 than when only Qk6 is depleted, indicating there are genes that may be regulated by both isoforms, or that respond in more complex ways to loss of both isoforms. QPCR measurement of expression changes for several genes from different classes validated this approach. Loss of Qk5 but not Qk6 causes *Hmga2* mRNA to go up while *Vcan* mRNA goes down, whereas the Qk6-specific gene *Fbn1* goes up as expected for a Qk6 responsive gene when either siQk5 or siQk6 is used for depletion (Fig S3E). Some genes like *Celf2* change expression upon Qk6 depletion but behave differently when both Qk5 and Qk6 are lost upon siQk5 treatment. Based on these data we conclude that each isoform has distinct roles in the expression of large, non-overlapping sets of genes, and that some genes may depend on both proteins in different ways for their expression.

We analyzed splicing using DEXSEQ and found that splicing changes are due primarily to loss of Qk5, as indicated from the reconstruction experiments in Fig 2A-C, and as observed previously (Hall et al. 2013). We compared the extent of splicing change in these samples to those generated by a pan-Qk siRNA analyzed on microarrays (Hall et al. 2013, Suppl. Table 3) and found these to be in good agreement (Fig S3F) confirming that Qk splicing functions are executed by Qk5.

We next asked whether Qk binding sites are enriched in the mRNA sequences of genes affected by loss of Qk by examining the distribution of CLIP peaks among genes with different responses to loss of Qk5 or Qk6 (Suppl Table 2H). Genes upregulated after loss of either Qk5 or Qk6 are significantly enriched for Qk binding sites (Fig S3G), suggesting that in many cases, Qk5 or Qk6 binding to mRNA represses mRNA levels. Genes upregulated after loss of Qk6 are not enriched, and those upregulated after loss of Qk5 are actually slightly depleted of Qk binding sites. This result suggests that at least some of the gene expression changes caused by loss of Qk5 or Qk6 may be direct, however others may be indirect, and could be mediated by other RNA binding proteins that Qk may control (Fig S3E, Zhao et al. 2010; Zearfoss et al. 2011; Mandler et al. 2014). Gene ontology analysis (Suppl. Table 2I) shows no enrichment of functional gene classes among Qk5-affected genes, however Qk6-affected genes are enriched for those involved in multicellular life. The extensive binding of Qk to its own 3’ UTRs (Fig 3E) suggests that Qk expression and control of appropriate amounts of the differently functioning Qk isoforms is regulated directly by Qk isoforms that bind isoform specific regions of each mRNA.

### Qk5 and Qk6 nuclear RNA accumulation requires Qk5 protein

Coverage tracks of RNA seq reads from siQk5, siQk6, and siNT libraries were examined to determine how siRNA depletion affected expression of Qk isoform mRNAs. Consistent with observation that loss of Qk5 results in loss of Qk6 protein, siQk5 causes reduction in the levels of spliced RNA for both Qk5 and Qk6 isoforms, whereas siQk6/7 treatment results in loss of only the Qk6-specific RNA (Fig 3E). This suggests that Qk5 is required for accumulation of RNA from the Qk locus, and explains the depletion of Qk6 protein after treatment with siRNA specific for Qk5. To validate this observation and to test whether the nuclear protein Qk5 is required for nuclear RNA accumulation, we fractionated siRNA treated cells and compared levels of Qk5 and Qk6/7 RNA in whole cells and in isolated nuclei (Figs 3F and G). As expected from the coverage tracks, siQk5 causes reduction of Qk5 and Qk6 total RNA, however siQk6 causes reduction of only Qk6 total RNA. Measurement of RNA from isolated nuclei (Fig S3H) shows that siQk5 reduces nuclear Qk5 and nuclear Qk6 RNA, whereas siQk6/7 has no effect on nuclear Qk5 RNA, but slightly (although not significantly) reduces nuclear Qk6 RNA (Fig 3G).

Qk5 might promote stability of Qk nuclear RNA by one or more of several mechanisms including most obviously splicing (Figs 2A, 2B, 3A, 3B), but also by promoting polyadenylation or other stabilizing events. To test this, we examined splicing at the Qk5 specific 3’ splice site for the last intron of the Qk5 mRNA, and evaluated read-through at the Qk5 poly(A) site relative to total nuclear Qk5 (Fig 3H). After treatment with siQk5, total Qk5 (Fig 3F) and nuclear Qk5 (Fig 3G) RNA is reduced, but the remaining RNA shows a relative increase in unspliced Qk5, suggesting that Qk5 promotes splicing of its own mRNA (Fig 3H). Depletion of Qk6 appears to activate splicing of Qk5, as the fraction of nuclear Qk5 that is unspliced is reduced upon loss of Qk6 (Fig 3H), indicating that one response of Qk6 depletion includes activating splicing of Qk5 mRNA, an effect consistent with other observations that loss of Qk6 promotes Qk5 through relief of translational repression by Qk6 (see below). There is no significant change in the fraction of RNA that is cleaved at the poly(A) site for Qk5 mRNA (Fig 3H).

To exclude the possibility that the dramatic RNA accumulation defect is a unique effect of siQk5-1, we tested two other Qk5 siRNAs that target distant regions of the Qk5 isoform mRNA, and found that each also reduces both Qk5 RNA and Qk6/7 RNA (Fig S3I). Absent a large contribution of this dramatic reduction by direct action of siQk5 on nuclear RNA, these results support a hierarchical model whereby initial production of small amounts of Qk5 promote increased expression of Qk5 mRNA through increased Qk5 specific 3’ splice site usage, leading to increased Qk5 protein, which then increasingly promotes Qk6 and Qk7 RNA expression.

### Testing isolated segments of the Qk gene to identify cross-isoform regulatory controls

Our attempts to understand Qk isoform regulation have been challenged by the very regulatory elements we sought to study, in particular the effect of endogenous Qk5 on assessment of Qk6 function (Figs 1 and 2), and the requirement of Qk5 for nuclear RNA production of Qk5 and Qk6 (Fig 3). To address this we have isolated segments of the Qk gene into reporters and studied expression of these segments after manipulation of levels of different isoforms (Fig 4A). First we wanted to confirm the effect of Qk5 loss on expression and splicing at the Qk5 specific 3’ splice site. We cloned this segment of the gene into the DUP reporter (Fig 4A center) and also created a version with a mutated 3’ splice site. Depletion of Qk5 (and Qk6/7) with siQk5 causes a reduction in both total RNA from the reporter gene (Fig4B left panel) as well as spliced Qk5 RNA (Fig4B right panel) relative to control siRNA, indicating that Qk5 is required for accumulation of Qk5 RNA. Mutation of the 3’ splice site abolishes the accumulation of spliced Qk5 RNA as expected, and leads to a nearly 50% reduction in the accumulation of RNA from the reporter, suggesting that splicing provides some but not all of the stabilization of RNA provided by Qk5. Since RNA from the reporter does not contain the target site for siQk5, it is unlikely that direct destabilization by siRNA explains this result. We conclude that Qk5 promotes accumulation of RNA from the Qk locus through splicing, as well as other unknown mechanisms.

**Figure 4:**
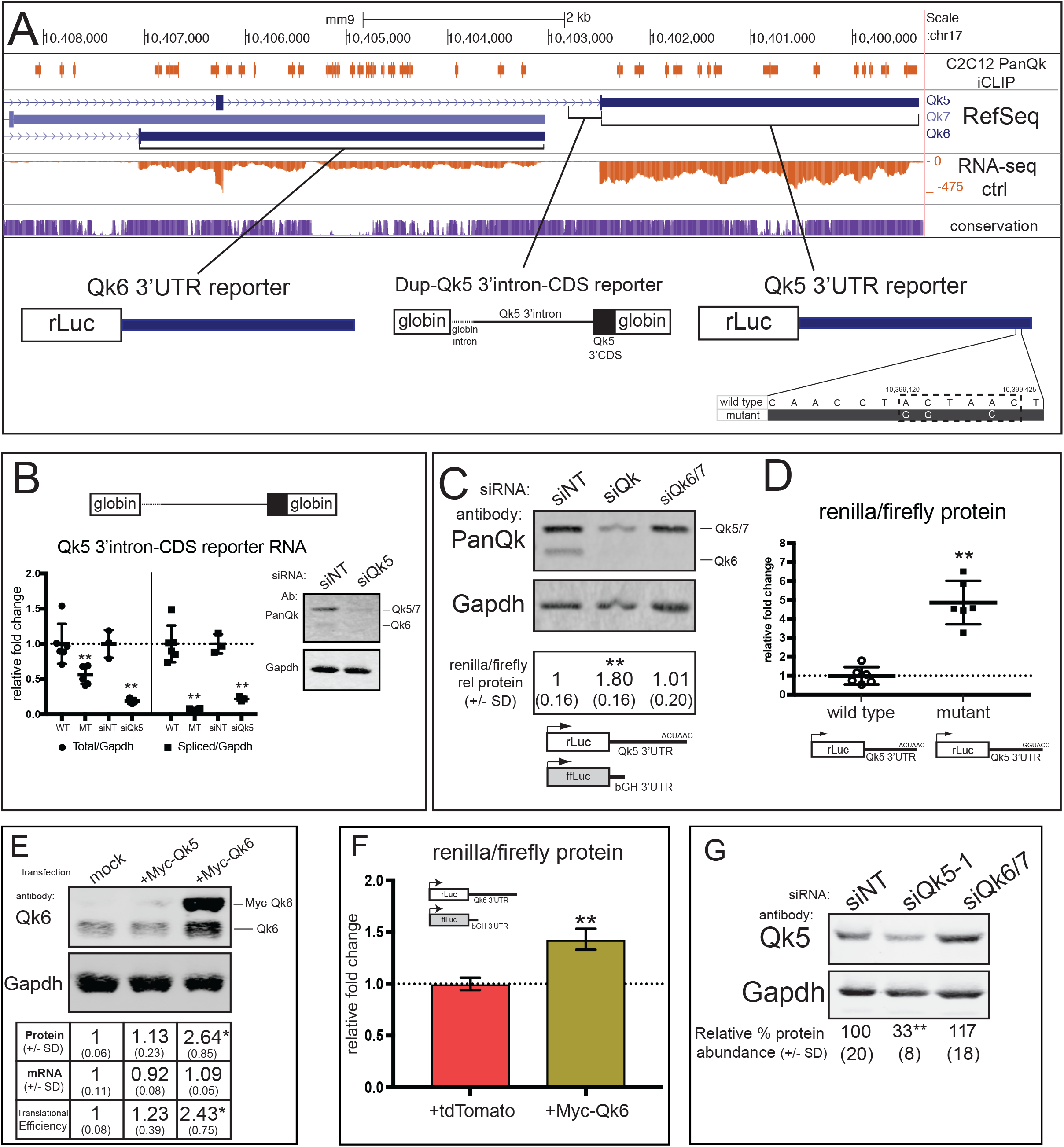
Auto- and cross-isoform regulation of Qk isoforms. A. UCSC Genome Browser screen shot showing the 3’end of Qk isoforms including PanQk iCLIP, RefSeq Genes, siNT RNA-seq coverage, and conservation tracks with regions of Qk sequence cloned into respective reporter gene constructs tested below. B. RT-qPCR performed on nuclear RNA extracted from C2C12 myoblasts transfected with Qk5 3’intron-CDS reporter (wild-type (WT) or 3’splice site mutant (MT)) or WT reporter plus siNT and siQk5-1 measuring total reporter RNA (left) and spliced reporter RNA (right) normalized to Gapdh RNA and reported on scatter plot as fold change relative to respective control (error bars represent standard deviation from the mean of 6 (WT vs. MT) or 3 (siNT vs. siQk5) biological replicates; ** p < 0.01). C. Western blot (top) of protein extracted from C2C12 myoblasts co-transfected with siNT, siQk, or siQk6/7 and renilla luciferase Qk5 3’UTR reporter and firefly luciferase, and renilla protein normalized to firefly protein relative to control siNT is shown below (mean values of 3 biological replicates shown +/- standard deviation (** p < 0.01). D. Scatter plot showing renilla/firely values of C2C12 cells transfected with either wild type or mutant Qk5 3’UTR reporter (n = 6; ** p < 0.01). E. Summary of endogenous Qk6 expression analysis during mock, Myc-Qk5, or Myc-Qk6 transfection of C2C12 myoblasts. Top shows western blot using Qk6 (top) and Gapdh (bottom) antibodies with observed migration pattern of Qk6 noted on right. Table below shows relative quantities of endogenous Qk6 protein (top, normalized to Gapdh), RNA (middle, normalized to Gapdh), and translational efficiency (bottom, relative protein/relative RNA; mean values of three independent replicates reported +/- standard deviation (* p < 0.05). F. Measurement of renilla/firefly protein from C2C12 myoblasts co-transfected with Qk6 3’UTR reporter and control tdTomato vector (left) or Myc-Qk6 (right); mean of 3 independent replicates +/- standard deviation reported (** p < 0.01). G. Western blot of whole cell protein extracted from C2C12 myoblasts transfected with siNT, siQk5-1 and siQk6/7 probed with Qk5 or Gapdh antibody; Qk5 protein level is normalized to Gapdh protein and mean percentage protein abundance relative to siNT calculated from 3 independent biological replicates is shown at bottom, +/- standard deviation.

### Qk5 inhibits its own expression by 3’UTR binding

We have provided evidence that Qk5 is required for *Qk* gene expression, especially when Qk5 levels are greatly reduced (Fig 3E-H, Fig 4A). Something must feed back to limit Qk5 expression when sufficiently high levels of Qk5 are reached. Previous studies of mammalian Qk5 (Larocque et al. 2002) or its homolog fly HowL (Nabel-Rosen et al. 2002) suggest a general role for Qk5 as a repressor of gene expression through binding to 3’UTRs, consistent with our CLIP enrichment results in which Qk5 repressed genes are enriched for Qk binding sites, but we also find enrichment of Qk6 repressed genes (Fig 3E, Suppl. Table 2H). One attractive site for such regulation occurs where we (Fig 3E, CLIP track) and others (Hafner et al. 2010; Van Nostrand et al. 2016) find Qk binding at a conserved computationally predicted binding site (Paz et al. 2014) near the end of the Qk5 3’ UTR. We cloned the entire Qk5 3’UTR into a renilla luciferase reporter and compared expression of this reporter to expression of a cotransfected firefly luciferase reporter carrying the bovine growth hormone 3’ UTR in cells depleted of either all Qk proteins (siQk) or Qk6 and Qk7 (siQk6/7, N.B. siQk5 targets this reporter and was not used). Depletion of all Qk protein forms, but not Qk6/7 alone, leads to a statistically significant 1.8 fold increase in renilla protein expression relative to firefly luciferase (Fig 4C). This suggests that Qk5 negatively regulates its own expression through its 3’UTR. To test whether the binding site near the end of the Qk5 3’UTR mediates this repression by Qk5, we made a reporter mutation to substitute UACUAAC (wild-type) with UGGUACC in the RNA (Fig 4A, right). This mutation leads to a nearly 5-fold increase in expression compared to wild type (Fig 4D). We previously noted that overexpression of Myc-Qk5 leads to a reduction in endogenous Qk5 mRNA and protein (Fig 4SA), so we tested whether overexpression of Myc-Qk5 would significantly downregulate the expression of the Qk5-3’UTR reporter, and it does (Fig S4B and S4C).

Qk proteins also bind the Qk6 mRNA 3’ UTR (Fig 3E(Hafner et al. 2010; Van Nostrand et al. 2016), and phylogenetically conserved Qk binding motifs are also found in this sequence (Paz et al. 2014). To test whether Qk5 similarly controls Qk6 through binding to the Qk6 3’ UTR, we made a reporter gene with the Qk6 3’UTR fused to the renilla luciferase coding region, and co-transfected it with a firefly luciferase control reporter, and either control, siQk, or siQk5-1 siRNAs (Fig S4D). We observe no change in reporter protein relative to control (Fig S4E) suggesting that Qk proteins do not repress expression through the Qk6 3’UTR reporter as they do through the Qk5 3’UTR.

Together these results suggest that Qk5 binds the UACUAAC element in its own 3’UTR to negatively regulate its expression. Thus, Qk5 protein at low levels promotes additional Qk5 mRNA accumulation through splicing and stabilization of nuclear RNA, but at high levels acts by binding its own 3’ UTR to limit expression through an as yet unknown mechanism. Our analysis of the distribution of Qk binding sites in genes specifically upregulated after loss of Qk5 indicates that many such genes have binding sites for Qk protein, suggesting that Qk5 mRNA is not the only mRNA whose expression is repressed by Qk5.

### Qk6 positively regulates its own translation through its 3’UTR

During overexpression experiments with Myc-Qk6 we noticed a reproducible increase in levels of endogenous Qk6. We repeated the overexpression of Myc-Qk6 in myoblasts and performed quantitative western blotting to measure both ectopic and endogenous Qk6 (Fig 4E). We observe a 2.5-fold increase in endogenous Qk6 protein without a significant RNA level change, indicating a 2.43-fold increase in translational efficiency (p < 0.05; Fig 4E). No significant change in endogenous Qk6 protein levels is observed when Myc-Qk5 is overexpressed, suggesting that increasing the level of Qk5 has little ability to further increase Qk6 RNA accumulation in these cells (Fig 4E). It is possible the slight and statistically insignificant increase in translation of Qk6 observed is due to increased levels of Qk5 in the cytoplasm, where it may contribute to normal Qk6 protein function by mislocalization.

To determine whether Qk6 autogenous control of protein abundance is mediated by translational control through its 3’UTR, we tested a luciferase reporter carrying the Qk6 3’UTR (Fig 4A, left), and compared its expression to the control firefly luciferase reporter with the bovine growth hormone 3’UTR after cotransfection with a myc-Qk6 expressing plasmid, or a control plasmid expressing tdTomato (Fig 4F, Fig S4F). Overexpression of Myc-Qk6 but not tdTomato protein leads to a significant increase in renilla expression mediated by the Qk6 3’UTR as compared to control (Fig 4F), indicating that the Qk6 3’UTR contains an autoregulatory element through which Qk6 activates its own translation. We conclude that Qk6 promotes translation of its own mRNA.

### Qk6 negatively regulates Qk5 protein expression

If Qk5 protein promotes expression of both its own and Qk6 mRNA (Fig 3), then reciprocal influences may exist whereby Qk6 protein feeds back to control Qk5 levels. We routinely observe a 15-25% increase in endogenous Qk5 protein levels when Qk6 is depleted using either of two Qk6 siRNAs (Figs 3C, Fig 4G, FigS4G and H). In other experiments where Qk6 is depleted, small but measurable effects in the direction opposite the effect of Qk5 depletion are often observed, such as statistically significant additional splicing repression (Rai14, Fig 3A) or activation (Capzb) of exons repressed or activated by Qk5, as would be expected when Qk5 levels increase. This increase in Qk5 protein level upon loss of Qk6 is accompanied by a consistent but modest reduction in Qk5 mRNA level relative to control (Fig 3F) to produce a 37% increase in translational efficiency (Vasudevan and Steitz 2007) of Qk5 mRNA when Qk6 is depleted. This opposing relationship between RNA stability and translational efficiency has been previously described (Kawai et al. 2004), and in this case suggests that Qk6 protein binding to the Qk5 mRNA stabilizes the transcript as it represses translation. Although these data are not statistically significant, they suggest that Qk6 represses translation of Qk5 mRNA, consistent with the function of Qk6 on other mRNAs in other cell types (Saccomanno L 1999; Zhao et al. 2010). The small effects of Qk6 depletion on endogenous Qk5 levels are not recapitulated using the Qk5 3’UTR reporter (Fig 4C) suggesting the possibility that sequences outside the 3’ UTR are required. We hypothesize that in cell types where Qk6 protein is more abundant, this regulation may predominate.

The results to this point identify several autoregulatory and cross-isoform regulatory controls that form a regulatory network that controls *Qk* expression and the composition of the different isoforms produced by the gene in myoblasts, in turn controlling large and distinct sets of other genes. Qk5 is the most abundant isoform and required for all *Qk* expression at the level of RNA accumulation (Fig 3F-H) and splicing (Fig 2), promoting its own and Qk6 expression at low levels (Fig 3F-H; Fig 4B), but also feeding back negatively on its own expression through a binding site in its 3’UTR (Fig 4C and D). Qk6 is less abundant but promotes its own translation (Fig 4E and F), while inhibiting Qk5 translation Fig 4G).

### Subtle inhibition of Qk translation reveals homeostatic responses of the Qk network

Evidence supporting the existence of the Qk isoform autoregulatory network has been obtained using siRNA depletions, designed to create catastrophic loss of protein, which may quickly overwhelm even a robust homeostatic network. In addition, siRNAs are subject to uncertainties about the extent to which they may target pre-mRNAs in the nucleus (Langlois et al. 2005; Berezhna et al. 2006), complicating interpretation. To test more subtle perturbations of the network and to avoid the use of siRNAs, we inhibited translation of all Qk mRNA using a morpholino oligonucleotide to occlude the translational start site shared by all Qk mRNAs, and evaluated the expression of Qk protein isoforms (Fig 5A) and RNAs in isolated chromatin and nucleplasmic fraction of nuclei and cytoplasmic fractions (Fig 5; (Pandya-Jones et al. 2013)). Treatment with the Qk start site morpholino reduced total expression of Qk by about 50% as compared to the control non-targeting morpholino (Fig 5A). Fractionation was monitored by following proteins (Fig 5A) and nuclear RNA distributions (Fig 5S). A panel of qRT-PCR primer pairs that span key regions of the Qk transcription unit are shown with their positions in the branched processing pathway with Qk6 specific RNAs on top and Qk5 specific RNAs at the bottom (Fig 5B, see also Fig 3E, Fig 4A). Upon subtle inhibition of Qk translation, the level of total Qk RNA increases relative to Gapdh RNA in all cell fractions tested (Fig 5C, left). This response is the opposite of that observed with catastrophic loss of Qk RNA observed with siQk or siQk5 (Fig 3). However, it is consistent with the observation that Qk5 represses its own expression via a Qk binding site near the end of its 3’UTR (Fig 4C and D). A homeostasis model would predict that lowering Qk5 protein would relieve Qk5 mediated repression of Qk5 resulting in an upregulation of Qk5 expression, which would increase Qk5 RNA levels in order to return to an appropriate level of Qk5 protein, after which repression would be re-established. Catastrophic loss of Qk5 mRNA may prevent this response because a threshold of functional Qk5 may be required to mount it.

**Figure 5:**
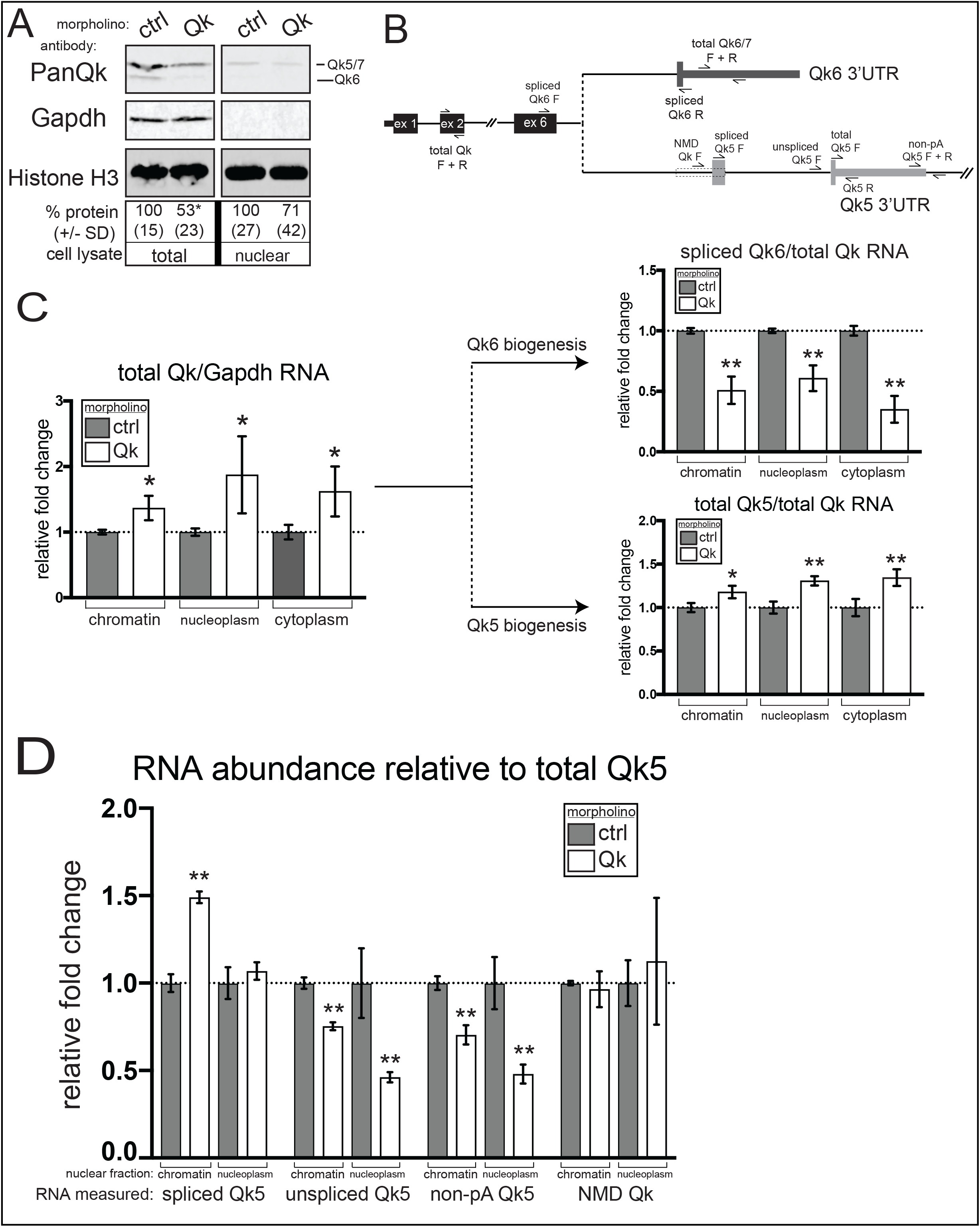
Moderate reduction of total Qk protein results in increased Qk5 RNA processing and decreased Qk6 RNA production. A. Representative western blot of whole cell protein extracts (left) or nuclear extract (right) from C2C12 myoblasts to which either control (ctrl) or Qk morpholino oligomer has been delivered simultaneously probed with PanQk, Gapdh, and histone H3 antibodies; quantitation of mean infrared signal for total Qk relative to Gapdh (left, total protein) or total Qk relative to histone H3 (right, nuclear fraction) reported below as percentage protein abundance relative to ctrl and +/- standard deviation of the mean calculated from 3 independent biological replicates (* p < 0.05). B. Schematic overview of Qk transcripts and location of qPCR primer sets used for subsequent measurements. C. RT-qPCR of RNA extracted from chromatin, nucleoplasmic, or cytoplasmic fractions of C2C12 myoblasts treated with ctrl or Qk morpholino measuring abundance of total Qk RNA normalized to Gapdh RNA (left), spliced Qk6 RNA normalized to total Qk RNA (top right), or total Qk5 RNA normalized to total Qk RNA (bottom right); all values reported as mean fold change relative to ctrl +/- standard deviation of the mean from 3 indpedent replicates (** p < 0.01; * p < 0.05). D. RT-qPCR of RNA extracted from chromatin or nucleoplasmic fractions of C2C12 myoblasts treated with ctrl or Qk morpholino measuring abundance of spliced Qk5 RNA, unspliced Qk5, polyadenylation read-through Qk5, or Qk NMD isoform each normalized to total Qk5 RNA; all values reported as mean fold change relative to ctrl +/- standard deviation of the mean from 3 indpedent replicates (** p < 0.01).

We next asked how inhibition of Qk translation affected the apportioning of total Qk transcripts in the directions of Qk5 mRNA (Fig 5C bottom right) and Qk6 mRNA production Fig 5C, top right). In all fractions tested, the relative amount of spliced Qk6 mRNA decreased significantly, whereas the relative amount of spliced Qk5 mRNA increased (Fig 5C, right). This response to subtle Qk protein depletion shows that alternative splicing has shifted in favor of the Qk5 isoform, a response that will lead to resetting of Qk protein and isoform composition by increasing Qk5 expression.

Finally we examined how morpholino inhibition altered the fraction of Qk5 processing precursors and products in the nucleus as a fraction of total Qk5 RNA (Fig 5 D). Spliced Qk5 (via the Qk5 specific last 3’ splice site, Fig 5B) associated with chromatin increases, concomitant with a decrease in unspliced Qk5 RNA. The change in amount of RNA that reads through the poly(A) site for Qk5 mRNA is reduced but mirrors the reduction in unspliced RNA (Fig 5D). We could detect no change in level of the NMD isoform of Qk5 in the nucleus. We conclude that the model for autoregulation of Qk5 explains the consequences of a subtle loss of Qk protein as follows: loss of repression of Qk5 on its own 3’UTR, followed by increased Qk5 expression that increases both overall accumulation of Qk RNA and directs splicing of more Qk5 mRNA. As Qk5 mRNA increases, Qk5 protein levels return and are sufficient to repress Qk5 again via its 3’ UTR. Qk6 levels return more slowly; as Qk5 levels reset, the splicing choice increasingly favors Qk6 mRNA, which is more efficiently translated as Qk6 protein levels are restored.

### Cross isoform Qk regulatory control is conserved in C6 glioma cells

Qk5 is the predominant Qk isoform present in C2C12 myoblasts (Fig 1B). In contrast, Qk6 and Qk7 are the predominant isoforms in adult mouse brain (Hardy et al. 1996). If the regulatory interactions we observe in C2C12 myoblasts are at play in other cells, then those with high Qk6 and low Qk5 might be explained as follows. The strong requirement of Qk5 for Qk6 RNA we observe in myoblasts might be less if the positive translational autoregulation of Qk6 (Fig 4G and F) were enhanced. Under such conditions, Qk6 might also more efficiently repress the translation of Qk5 mRNA (Fig 4H). To begin testing this network in other cells, we used quantitative western blotting) to measure Qk5, Qk6, and Qk7 protein abundance (Fig 6A and B). C2C12 cells have a high Qk5/Qk6 ratio (with Qk7 nearly undetectable; Fig 1A; Fig 6A and B). Consistent with previous findings (Hardy et al. 1996), adult mouse optic nerve and cerebellum tissue have distinct Qk5/Qk6/Qk7 compositions, and a much lower Qk5/Qk6 ratio (Fig 6A). In contrast, rat C6 glioma (Benda et al. 1971) and CG4 oligodendrocyte precursor cells (Sharma et al. 2011) have intermediate Qk5/Qk6 ratios, with different amounts of Qk6 or Qk7 (Fig 6A and B). We chose rat C6 glioma cells as a counter example to mouse C2C12 cells to test whether the autoregulatory controls observed above operate in cells with lower Qk5/Qk6 ratios.

**Figure 6:**
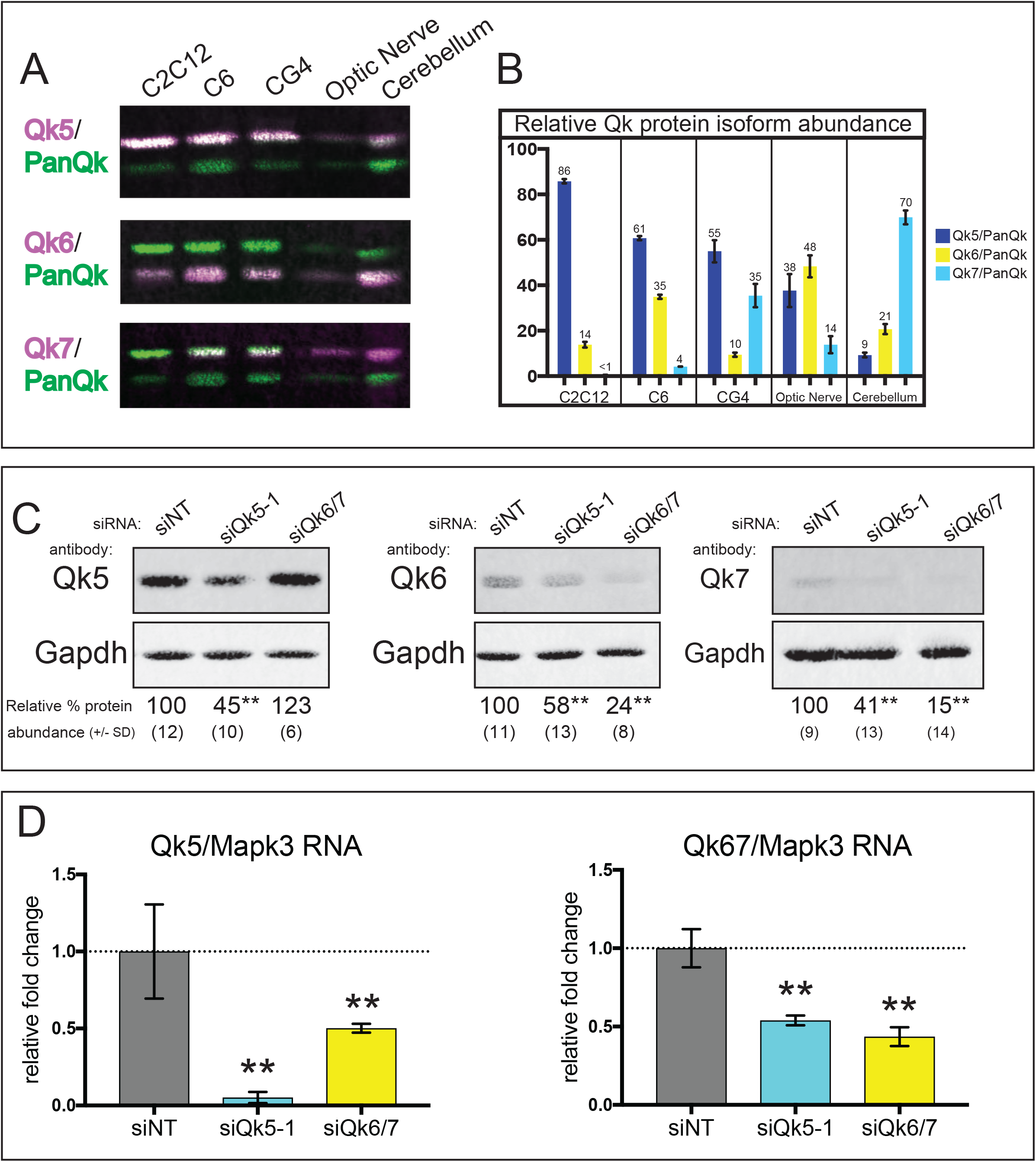
Qk cross-isoform regulation is conserved in rat C6 glioma cells. A. Representative western blots of whole cell protein extracts from C2C12 myoblasts, rat C6 glioma cells, rat CG4 oligodendrocyte precursor cells, mouse optic nerve tissue, and mouse cerebellum tissue probed with Qk5 (top), Qk6 (middle), or Qk7 (bottom) in magenta channel and PanQk in green channel B. Western blots (see 6A) infrared intensity values for each isoform were normalized to PanQk and used to determine mean percentage of each protein isoform relative to total (PanQk) in whole cell protein extracts from C2C12 myoblasts, rat C6 glioma cells, rat CG4 oligodendrocyte precursor cells, mouse optic nerve tissue, and mouse cerebellum sampled in biological triplicate. C. Representative western blots of whole cell protein extracts from rat C6 glioma cells transfected with siNT, siQk5-1, and siQk6/7 simultaneously probed with Qk5 and Gapdh (left), Qk6 and Gapdh (middle), and Qk7 and Gapdh antibodies; quantitation of mean infrared signal for Qk isoform relative to Gapdh reported below as percentage protein abundance relative to siNT and +/- standard deviation of the mean calculated from 3 independent biological replicates (** p < 0.01). D. RT-qPCR of RNA extracted from rat C6 glioma cells described in 6C measuring Qk5 RNA (left) or Qk6/7 RNA (right) normalized to Mapk3 mRNA and displayed as fold change relative to siNT +/- standard deviation (** p < 0.01).

We treated C6 glioma cells to deplete either the Qk5 or Qk6/7 isoforms (the siRNA target sequences are conserved in mouse and rat). Although overall siRNA-mediated depletion appears less efficient in these cells as compared to C2C12 cells, the loss of Qk6/7 upon depletion of Qk5 is evident (Fig 6C), indicating that the hierarchical requirement of Qk5 for expression of Qk6/7 RNA (Fig 6C) and protein (Fig 6B) is conserved. The magnitude of the reduction of Qk6 protein in C6 glioma cells is less than that observed in C2C12 myoblasts (Fig 6C compare to Fig S3A). We also observe an increase in Qk5 protein under Qk6/7 depletion in C6 glioma cells (Fig 6D), indicating that the feedback repression of Qk6 on Qk5 (Fig 3G) is conserved. In contrast to C2C12 cells where siRNA depletion of Qk6 does not affect Qk5 mRNA, C6 cells suffer a significant reduction of Qk5 RNA under this condition (Fig 6D), which represents a 2.6-fold increase in translational efficiency in C6 glioma cells (p < 0.01) compared to a (nonsignificant) ~1.4-fold in C2C12 myoblasts upon loss of Qk6/7. Due to the higher relative concentration of Qk6 in C6 glioma cells, the translational repression of Qk5 is more efficient. Taken together, these findings suggest that the *Qk* autogenous regulatory network operates at various settings in different cells (Fig 7A), where the Qk5/Qk6/Qk7 composition can vary in a stable way, using parallel sets of auto- and cross regulatory control. The existence of multiple different stable settings of the Qk5/Qk6/Qk7 composition indicates that the *Qk* regulatory network is responsive to the influence of other regulatory proteins and RNAs, which may add controls on top of the network to achieve homeostasis at different Qk5/Qk6/Qk7 compositions.

**Figure 7:**
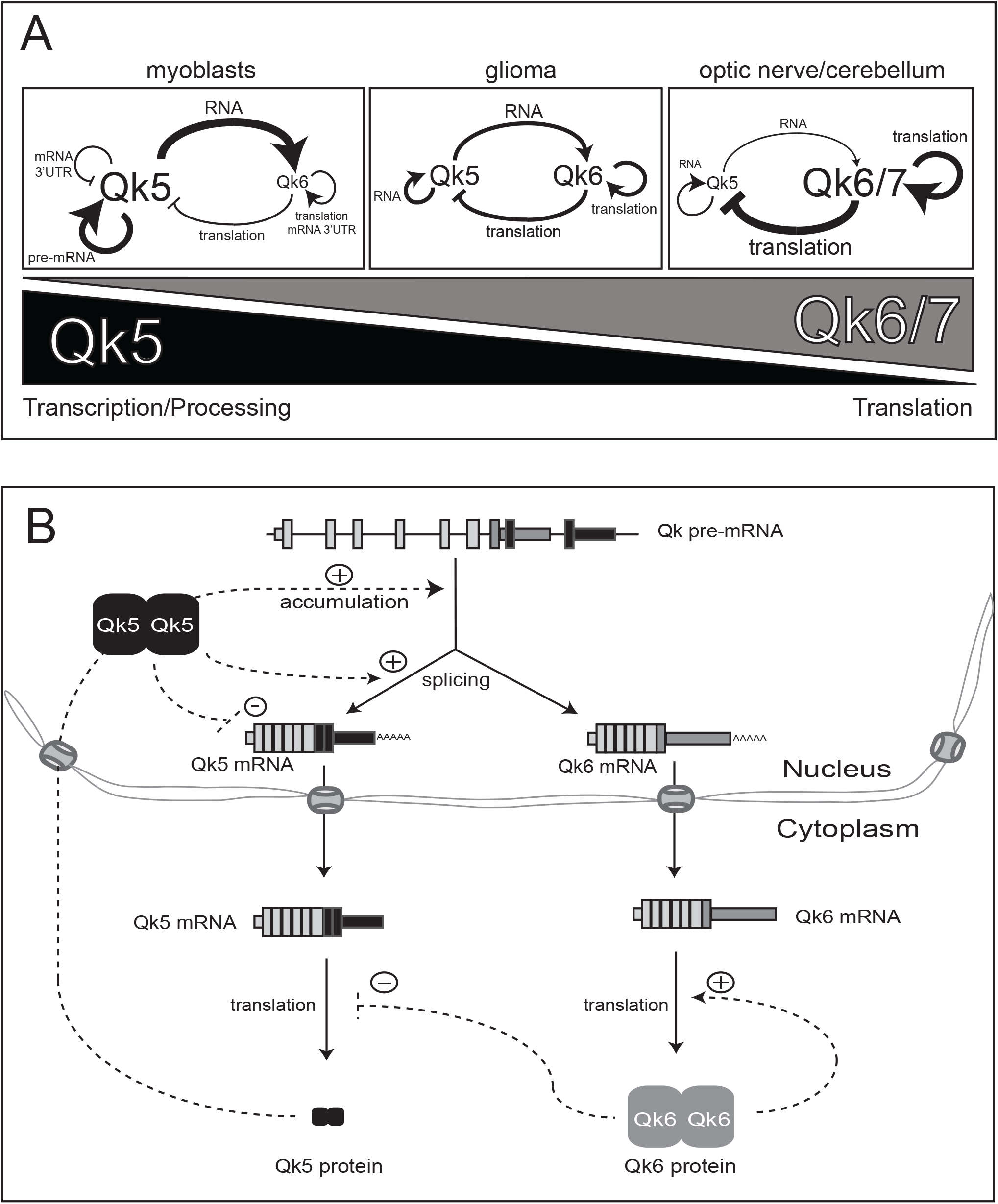
Models of Qk autogenous regulation. A. Quantitative rheostat model: various relative levels of Qk protein isoforms are observed in different cell/tissue types such as Qk5 high/Qk67 low in C2C12 cells, medium Qk5/medium Qk67 in rat C6 glioma cells, and high Qk67/low Qk5 in adult mouse optic nerve and cerebellum tissues. These different relative protein levels support the observed auto- and cross-isoform regulatory interactions defined in this study, albeit by imposing greater influence (thicker lines) on specific processing steps when their relative concentration is higher (larger font) than in other cell/tissue types. Arrows denote positive regulation while lines ending in perpendicular line denote negative regulation. B. Hierarchical model: the *Qk* gene transcribes a single pre-mRNA which is processed to make mature Qk mRNA isoforms which are then exported from the nucleus and translated into Qk5 and Qk6 proteins. The NLS unique to the Qk5 C-terminus mediates nuclear import, thus more Qk5 protein is observed in the nucleus. Here Qk5 promotes nuclear accumulation of Qk pre-mRNA and formation of Qk5 mRNA through a splicing based mechanism. Accumulation of Qk5 protein negatively feeds back to reduce its own levels, while it also positively feeds back to promote accumulation of Qk6 mRNA. Qk6 protein is predominantly localized in the cytoplasm and negatively regulates the translation of Qk5, and positively regulates its own translation. Dotted lines represent regulatory interactions, while solid lines represent spatial and temporal flow of genetic information.

## Discussion

We initiated this study to understand how RBP family isoforms that recognize the same RNA sequence in different functional contexts (splicing, RNA transport, translation, decay) are regulated within a single cell so that each of these processes is appropriately supplied with the correct amount of RBP. We chose Qk because all isoforms arise form a single gene, to avoid complexities that exist with other RBP families that expressed multiple isoforms from each of multiple genes. In the case of *Qk*, isoform function is largely determined by cellular compartment: nuclear functions (splicing, nuclear RNA accumulation) are executed by the predominantly nuclear isoform Qk5, whereas cytoplasmic functions (translational control, mRNA decay) are executed by the predominantly cytoplasmic Qk6 (Figs 1, 2, and 3 (Hardy et al. 1996; Lu et al. 2003)). We note that both isoforms appear to have dynamic subpopulations outside their predominant localization sites, and that the two-dimensional imaging method we used may overestimate nuclear localization due to cytoplasm above and below the nucleus (Fig 1). Homeostasis and control of the levels of each isoform is established through a network of auto- and cross regulatory controls (Fig 7) whereby Qk5 is required for its own and Qk6 RNA expression from the nucleus (Fig 3), and Qk6 activates its own and represses Qk5 mRNA at the translational level (Fig 4). The strength of these controls is tunable such that cells express different ratios of Qk5 and Qk6 in a stable characteristic way (Fig 7).

### A division of labor between Qk protein isoforms

We previously identified Qk protein as a regulator of muscle-specific alternative splicing through binding to intronic ACUAA sequence motifs (Hall et al. 2013). Since the Qk5 isoform contains a NLS (Wu et al. 1999), we expected Qk5 to be responsible for splicing. Indeed, Qk5 is both necessary and sufficient for splicing regulation (Figs 2 and 3), whereas Qk6 and Qk7 are dispensable (Fig 2). Although artificially overexpressed Myc-Qk6 can activate splicing (Fig 1F), abnormally high nuclear localization after increased expression in cells with already high levels of Qk5 is likely a consequence of heterodimerization with endogenous Qk5 that increases nuclear concentration of Qk dimers (Fig 1F, (Pilotte et al. 2001). A mutant Qk protein containing all the common Qk sequences but lacking isoform-specific C-termini (QkBody) is distributed throughout the nucleus and cytoplasm, and suffices for splicing, whereas Qk6 does not (Fig 2). Careful comparison of the cellular distribution of the QkBody protein with that Qk6 indicates that the 8 Qk6-specific C-terminal amino acids encode a cytoplasmic retention or nuclear export signal (Nakielny and Dreyfuss 1999; Cook and Conti 2010) that increases its cytoplasmic representation as compared to QkBody (Fig 2G and H). Qk5 appears to control events other than splicing, by unknown mechanisms that lead to nuclear RNA accumulation (Fig 3F-H, Fig 4B) at least a part of which seems independent of splicing per se (Fig 4B). Qk5 also represses its own expression through a Qk binding site in its 3’UTR, possibly through nuclear retention (Larocque et al. 2002; Nabel-Rosen et al. 2002), but the mechanism and subcellular location of this repression is unknown. Qk6 protein is an activator of its own translation (Figs 4E and 4F), a function consistent with its predominantly cytoplasmic localization (Zhao et al. 2010). Since these distinct functions correspond with the main subcellular distributions of each isoform, and isoform specific C-terminal protein sequences seem dedicated only to localization, we suspect that isoform functions are enforced primarily by localization. It follows that all Qk protein elements required to interact with the splicing, translation, export and decay machineries lie within QkBody, and that additional dissection of Qk protein sequences will be necessary to map the sites required for function in each of those processes.

Our analysis of the broader effects of loss of specific Qk isoforms indicates that the specific regulatory functions identified in reporter tests extend into the transcriptome. We identified sets of mRNAs whose levels significantly increase or decrease upon loss of Qk5 or Qk6 (Figs S3C-E, Supplemental Tables 2A-I). In particular, distinct sets of mRNAs increase in level after loss of Qk5 or Qk6 are enriched for Qk binding sites identified by CLIP (Fig 3D, Fig S3G), similar to the Qk5 mRNA itself (Fig 4C and D). The existence of separate sets of mRNAs whose levels are controlled by only one Qk isoform or the other is further support for the idea that the division of labor between these isoforms, enforced by distinct subcellular localization, is deeply integrated into cell function.

### Auto- and cross isoform regulation controls Qk isoform composition

Our initial effort to deplete cells of each single isoform of Qk protein in turn was complicated by the strong dependence of Qk6 expression on expression of Qk5 (Fig 3). We were able to construct cells depleted of all endogenous isoforms and replaced with a single isoform to confirm that Qk5 is responsible for splicing (Fig 2A-C). Furthermore the Qk5 isoform-specific C-terminal tail containing the non-canonical NLS does not contain an essential splicing-specific function: it can be replaced by an N-terminal SV40NLS to promote splicing. In addition it seems that any manipulation that increases the amount of the common Qk sequences in the nucleus including recruitment of Qk6 or QkBody by heterodimerization with Qk5 will help activate splicing (Fig 1F-G, Fig 2D-F). In the remaining experiments, using such depleted and reconstructed cells was impractical, and we assigned a function to Qk5 if Qk5 depletion compromised that function but Qk6 depletion did not. If depletion of both Qk5 and Qk6 compromised a function, we tentatively assigned that function to Qk6, because depletion of Qk5 leads indirectly but efficiently to depletion of Qk6. It is important to note the danger in this conclusion, because our transcriptome analysis identifies genes whose expression does not fall neatly into two categories, probably because like Qk, they are regulated by both Qk isoforms to different extents.

In analyzing the effect of depleting each Qk isoform on the expression of RNA and protein for itself and the other Qk isoforms using a combination of endogenous Qk expression (Fig 3) and artificial reporter constructs (Fig 4), we found that Qk5 is required for nuclear accumulation of both its own transcripts and those encoding Qk6/7 (Fig 3). We also provide evidence that Qk6 represses translation of Qk5 mRNA but activates translation of its own mRNA (Fig 4). These results allowed us to build a model of Qk autogenous and cross-isoform regulation for C2C12 cells (Fig 7). This control network is more similar to a rheostat than a bistable switch where either Qk5 or Qk6 expression dominates, because it supports stable intermediate isoform mixtures from the *Qk* gene in different cell types (Fig 7A). The structure of this network is such that the same compartmentalization that constrains the functional role of each isoform is also used to create the auto- and cross-regulatory controls. Qk5 acts on its own RNA and that of Qk6/7 through nuclear processes, whereas Qk6 acts on both its own mRNA and that of Qk5 through translation and decay in the cytoplasm. This network controls both the total amount of Qk protein by combining a strong dependence of Qk6 expression on Qk5 with negative feedback of Qk5 by both Qk5 (at the RNA level) and Qk6 (at the translational level). Autoregulation by RBPs is common and can be imposed at different RNA processing steps including splicing (Wollerton et al. 2004; Lareau et al. 2007; Ni et al. 2007), polyadenylation (Dai et al. 2012), mRNA stability/decay (Ayala et al. 2011), and translation (de Melo Neto et al. 1995; Wu and Bag 1998). Although cross-paralog regulation exists between RBP family members encoded by separate genes (Boutz PL 2007b; Spellman et al. 2007; Wang et al. 2012)), the Qk regulatory network highlights the role of compartmentalization in both functional specification and role in the regulatory network of RBP isoforms (Fig 7). Other RBP families may use similar mechanisms for autogenous and cross-isoform control but for those encoded by multiple genes, additional feedback on transcription are possible and may be necessary.

Having described this network using siRNA depletion strategies, we were concerned about two potentially confounding effects. One is that catastrophic loss of mRNA for Qk may compromise the ability of the cell to respond sufficiently to restore homeostasis. We tested the potential of the network to produce a homeostatic response by an orthogonal method of inhibiting expression, by using a morpholino to block the translation start site sequences shared by all Qk isoform mRNAs (Fig 5). This showed that instead of complete loss of nuclear RNA observed with siRNAs against Qk5 mRNA, translational inhibition allowed cells to respond by increasing RNA levels from the Qk gene (Fig 5C). This could be due to promotion of nuclear RNA accumulation by Qk5, or could be an indirect effect on the rate of *Qk* transcription, but in either case Qk5 mRNA increases, as does Qk5 splicing at the expense of Qk6 splicing, as a fraction of total RNA from the *Qk* locus. This provides strong evidence that Qk5 helps control the ratio of Qk5 and Qk6 mRNA from the gene through alternative splicing.

Since the quantitative output from the Qk network appears to be set at different Qk5/Qk6/Qk7 ratios in different cell types (Fig 6A), it must also be sensitive to regulation by other factors. For example, the RBP RBFOX2 reduces Qk7 expression by repressing alternative splicing in human embryonic stem cells (Yeo et al. 2009). Since RbFox2 is an abundant splicing regulator in C2C12 cells (Bland et al. 2010; Singh et al. 2014), this may explain why Qk7 is nearly undetectable (Figs 1B and 6A). However, this seems not to hold in mouse cerebellum, where both Qk7 (Fig 6A) and RbFox2 (Gehman et al. 2012) are abundant. In another example, miR-214-3p targets regions common to both the Qk6 and Qk7 3’UTR (van Mil et al. 2012) which may specifically reduce Qk6 and Qk7 (Irie et al. 2016) or reduce total Qk (Shu et al. 2017). There likely are multiple additional means by which Qk isoform ratios can be regulated, but the Qk auto- and cross-regulatory isoform network uncovered here represents the foundational structure on which that regulation is imposed.

### Qk isoform ratio regulation, development, and cancer

*Qk* is a key regulator of development (Ebersole et al. 1996; Li et al. 2003; Justice and Hirschi 2010) and cancer (Novikov et al. 2011; Chen et al. 2012; Zong et al. 2014). Our model predicts that Qk5 expression precedes expression of other Qk isoforms temporally as transcription is upregulated at the *Qk* gene (Fig 7B) in a developmental context. This expectation appears to hold for embryogenesis in *Xenopus* (Zorn et al. 1997), zebrafish (Radomska et al. 2016), and mouse (Ebersole et al. 1996; Hardy et al. 1996). Additionally, early stem/progenitor cells in *Drosophila* (Nabel-Rosen et al. 1999) and mice (Hardy 1998), express predominantly Qk5 (or its ortholog HowL in *Drosophila*). In mice, *qk* knockout is embryonic lethal (Li et al. 2003) and the *qk*^*l1*^ mutant mouse, which appears to lack Qk5 expression due to disruption of Qk5-specific splicing by a point mutation (Cox et al. 1999), is recessive lethal (Shedlovsky et al. 1988). In this mutant, both Qk6 and Qk7 mRNAs are significantly reduced in visceral endoderm dissected from non-viable embryos (Cox et al. 1999; Bohnsack et al. 2006), suggesting that Qk5 is required for efficient Qk6 and Qk7 expression in this tissue also. These observations provide additional evidence in support of our model of *Qk* autoregulation, and suggest its importance in an evolutionarily conserved developmental context.

The extensively studied *qk^v^* mouse model of dysmyelination (Sidman et al. 1964) expresses reduced mRNA levels of all Qk isoforms in glia (Lu et al. 2003) and vascular smooth muscle (van der Veer et al. 2013), due to a deletion that includes a portion of the *Qk* promoter (Ebersole et al. 1996). Based on the models we present here (Fig 7), lower Qk5 protein levels in *qk*^*v*^ oligodendrocyte precursor cells may be insufficient for robust Qk6 and Qk7 up-regulation that normally occurs around the peak of myelination (Ebersole et al. 1996). Since deletion of *qk* in oligodendrocytes is lethal (Darbelli et al. 2016), the reduction in Qk6 and Qk7 in *qk*^*v*^ oligodendrocytes could be due to selective pressure imposed on Qk5 substrates to maintain viability of the organism. Therefore, the imbalance of Qk isoforms in *qk*^*v*^ oligodendrocytes results in developmental defects and the tremor phenotype without sacrificing viability of the organism.

Finally, disruption of tissue-specific Qk isoform expression patterns is also observed in non-small-cell lung cancer (NSCLC) (de Miguel et al. 2016; Sebestyen et al. 2016) and glioblastoma (Jin et al. 2004) patient samples (Human Qk is encoded by the *QKI* gene). Although overall Qk protein expression is reduced, an isoform ratio switch occurs whereby healthy lung cells expressing mostly Qk6 transition to a state in NSCLC samples where cells express predominantly Qk5 (de Miguel et al. 2016; Sebestyen et al. 2016). This isoform switch likely shifts *QKI* function away from cytoplasmic regulation such as translation and decay (Saccomanno L 1999; Zhao et al. 2010), toward nuclear processes such as splicing and primary transcript stability. Understanding the relationship of *QKI* gene function in the context of cancer and other diseases requires evaluation of both the specific functions of Qk protein isoforms on their own RNAs as well as on the other pre-mRNAs and mRNAs from the many other genes that *QKI* regulates.

## Materials and Methods

### Cell Culture

C2C12 cells were routinely cultured in Dulbecco’s modified Eagle medium (DMEM) with high glucose (Life Technologies) supplemented with 10% heat inactivated fetal bovine serum (Life Technologies) at 37°C with 5% CO2. For differentiation experiments, C2C12 cells were allowed to reach ~90% confluency, then media was changed to DMEM supplemented with 5% horse serum (Life Technologies) and the cells were harvested 72h post media change. C6 glioma cells were cultured in F-12K media (ATCC) supplemented with fetal bovine serum at 2.5% and horse serum at 15% as described (Benda et al. 1968) at 37°C with 5% CO2. CG4 oligodendrocyte precursor cells were grown in DMEM with high glucose supplemented with N1 (5 µg/ml transferrin, 100 µM putrescine, 20 nM progesterone, and 20 nM selenium), biotin (10 ng/ml), insulin (5 µg/ml), and 30% B104 cell conditioned media as described (Sharma et al. 2011) at 37°C with 5% CO2; all supplements purchased from Sigma-Aldrich.

Additional methods and oligonucleotide sequences found in Supplemental Information. RNAseq and CLIPseq reads analyzed in this paper can be found under GEO accession number GSE102615.

## Acknowledgements

We thank Sean Ryder, Doug Black, Jeremy Sanford, Mariano-Garcia Blanco, Shelton Bradrick, Rhonda Perriman, David Feldheim, Jason Talkish, and Susan Strome for reagents, discussion, and advice. This work was supported by National Institutes of Health (NIH) grant R01-GM040478 to M.A., NIH pre-doctoral T32 GM008646 to WSF, and California Institute of Regenerative Medicine (CIRM) pre-doctoral fellowship award TG2-01157 to WSF.

